# Convolutional neural networks can identify brain interactions involved in decoding spatial auditory attention

**DOI:** 10.1101/2023.11.22.566213

**Authors:** Keyvan Mahjoory, Andreas Bahmer, Molly J. Henry

**Author notes:** Corresponding authors (KM), (MH).

## Abstract

Human listeners have the ability to direct their attention to a single speaker in a multi-talker environment. The neural correlates of selective attention can be decoded from a single trial of electroencephalography (EEG) data. In this study, leveraging the source-reconstructed and anatomically-resolved EEG data as inputs, we sought to employ CNN as an interpretable model to uncover task-specific interactions between brain regions, rather than simply to utilize it as a black box decoder. To this end, our CNN model was specifically designed to learn pairwise interaction representations for 10 cortical regions from five-second input trials. By exclusively utilizing these features for decoding, our model was able to attain a median accuracy of 77.56% for within-participant and 65.14% for cross-participant classification. Through ablation analysis together with dissecting the features of the models and applying cluster analysis, we were able to discern the presence of alpha-band-dominated inter-hemisphere interactions, as well as alpha-and beta-band dominant interactions that were either hemisphere-specific or were characterized by a contrasting pattern between the right and left hemispheres. These interactions were more pronounced in parietal and central regions for within-participant decoding, but in parietal, central, and partly frontal regions for cross-participant decoding. These findings demonstrate that our CNN model can effectively utilize features known to be important in auditory attention tasks and suggest that the application of domain knowledge inspired CNNs on source-reconstructed EEG data can offer a novel computational framework for studying task-relevant brain interactions.

## Introduction

In a competing talker situation with noise, a healthy human can focus on a single talker. It has been shown that this focus is reflected in neural activity that more consistently tracks the temporal dynamics of the attended talker’s speech compared to the unattended talker’s speech [1,2]. Auditory selective attention abilities may be weakened or lost as a result of normal aging or hearing impairment. A promising way to potentially counteract selective attention impairment involves the automatic detection of the focus of auditory attention from neural activity and the subsequent amplification of the corresponding audio stream by hearing prostheses [3]. Most studies typically focus on decoding auditory attentional focus using EEG recordings as a non-invasive, portable, and less costly technique as opposed to magnetoencephalography (MEG) or intracranial EEG.

In the past, various approaches have been proposed to decode auditory attention from neural time series. Building on the observation that neural tracking of the amplitude envelope of speech is stronger for attended than unattended material, some studies have attempted to reconstruct the envelope of the attended speech from the EEG signal using linear models [2,4], state-space based models [5,6], canonical correlation analysis [7], and artificial neural networks [8–10]. Other studies have focused on decoding the spatial locus of auditory attention rather than the envelope of the attended speaker [11–14]. Studies focusing on decoding the spatial locus of auditory attention have revealed the importance of activity originating from a frontoparietal network in decoding accuracy [15,16]. In particular, it has been shown that the alpha activity originating from parietal areas is lateralized during selective attention to a location in space [17–20], and this lateralization can be leveraged to decode auditory spatial attention. A recent study has revealed the presence of at least two distinct generators of alpha oscillations over central and parieto-occipital areas during spatial auditory attention [17,21,22]. In one study, frontal beta-band activity was shown to be the main predictor of spatial auditory attention [14].

In the last decade, Deep Learning (DL) has emerged as the method of choice for a variety of tasks in computer vision, natural language processing, and audio recognition [23,24]. However, its applications to neural signals have encountered challenges due to certain characteristic properties of neural signals that distinguish them from image or audio data. For example, neural time series possess nonstationary temporal dynamics and spatial patterns occurring in specific frequency bands but typically with a low signal-to-noise ratio. In addition, EEG recordings contain measurement artifacts like eye movements, heart artifacts, and other unwanted noise sources. These properties substantially change the approach to training artificial neural networks for EEG signal decoding.

Despite the existing challenges, DL has recently demonstrated promise in helping make sense of neurophysiological signals [25,26]. Among several DL techniques, convolutional neural networks (CNNs) have been applied with some success to EEG classification tasks. Indeed, since 2015, CNNs have been the most common architecture type in the majority of EEG studies and their application has been growing steadily [25,27]. This amount of interest in CNNs has been attributed to their innate capability in end-to-end learning and their capacity in extracting temporal and spatial structures in the EEG data, as well as their successful applications on computer vision tasks [27,28]. For example, CNNs have been used for decoding auditory attention [14,29], seizure prediction and detection [30,31], and sleep stage classification [32–34].

Some studies have attempted to adapt existing CNN architectures to the task of decoding EEG data specifically, rather than importing them directly from the computer vision applications without modification. These updates in the CNNs architecture have enabled them to learn neurophysiologically interpretable features. For example, Schirrmeister and colleagues introduced a CNN architecture designed that can adapt spatial and temporal filters in the convolutional layers [27]. Along the same lines, Lawhern and colleagues proposed EEGNet, a CNN model adapted to the temporal and spatial properties of EEG data with a relatively small number of parameters to fit, that outperformed other models on four different data sets [28]. In the following years, this type of CNN architecture, built upon learning temporal and spatial features from EEG, has been an active field of research [35–37].

Sensor-level EEG recorded from the scalp, behind the ear, or ear canal, is an obvious choice for real-time applications e.g., the development of neuro-steered hearing aids, because of its portability, low cost, and noninvasive nature. However, EEG captures mixtures of neural activity originating from the entire brain, and not necessarily from the brain area beneath the electrodes [38]. This mixing poses challenges in making interpretations about brain regions based on sensor space EEG. Thus, sensor EEG may be suboptimal in the case that we want to use CNNs to make inferences about the underlying brain regions engaged in auditory decoding.

For this reason, in the current study, we utilized source-reconstructed EEG data together with a new CNN architecture and attempted to use CNNs as a data-driven analysis tool to identify cortical areas and their interactions in decoding spatial auditory attention. For this purpose, we used EEG data, recorded from 18 participants attending a speech stream presented to one ear and ignoring speech presented to the other. We reconstructed source time courses for 10 cortical regions including the left and the right occipital, temporal, parietal, central, and frontal areas. Time courses of the 10 cortical regions were used as input to our CNN model. In addition, the architecture of our CNN model was specifically designed to enable it to learn internally the interactions between the 10 cortical regions relevant to auditory attention. Our findings demonstrated that the trained CNN model makes use of features that are well-known to be essential for decoding auditory attention.

## Results

### Decoding performance

To decode auditory spatial attention based on source-reconstructed EEG, we used our CNN model, which due to its specific architecture, is able to capture interactions between 10 predefined cortical areas and utilize these features for decoding. Panel A of Figure 1 illustrates the architecture of our model. The model input comprises matrices of 30 x T dimensions, where 30 denotes the signals collected from 10 brain regions, and T represents the number of time points within the sliding window. The signals have undergone a frequency band specific dimensionality reduction to enhance activity within the delta-theta, alpha, and beta frequency bands. Input data were generated by moving a sliding window of size T with 50% of overlap over time points. We tested four values for T (1 s, 2 s, 5 s, and 10 s). Panel B of Figure 1 shows the decoding accuracy of our CNN model for four different lengths of input data (1 s, 2 s, 5 s, and 10 s), obtained from two training approaches: within-participant (left panel) and cross-participant (right panel). The within-participant decoder for a participant was trained, validated, and tested solely on the data of that participant, using a block-wise four-fold cross validation approach. In contrast, the cross-participant classifier for a participant was trained on data of other participants and then tested on that participant’s data to obtain a participant-independent model which generalizes across participants (see method section for further details).

Next, to explore the impact of input length on decoding accuracy, we used a Linear Mixed Effects Model (LMEM), where we specified the length of input data as the independent variable, the decoding accuracy as the dependent variable, and participants as a random effect. Our LMEM analysis, using the Statsmodels library (https://www.statsmodels.org/), found a significant effect of input length on decoding accuracy (within-participant: p < 0.001, cross-participant: p < 0.001). Furthermore, we found a significant increase in median decoding accuracy when the input length increased from 2 s to 5 s (Wilcoxon matched-pairs signed rank test, within-participant: W = 2, p < 0.001, cross-participant: W = 1, p < 0.001). We found a marginally significant improvement in decoding accuracy when the data input length was increased from 5s to 10s for within-and cross-participant decoding (within-participant: W = 39, p = 0.044; cross-participant: W = 37, p = 0.034). We did not observe any significant improvement in decoding performance when input data were increased from 1 s to 2 s for either decoding approach (within-participant: W = 53, p = 0.17; cross-participant: W = 44, p = 0.07). Thus, for the subsequent analyses, we chose the input samples generated by a sliding window of 5 s with 50% overlap, as a good trade-off between the length of input data and the performance of model for the classification task of decoding right from left auditory attention.

Panel C of Figure 1 compares the performance of our CNN model between the within-participant and cross-participant decoding approaches. Unsurprisingly, within-participant classification (median = 77.56) outperformed cross-participant classification (median = 65.14). For the majority of participants (88%), our CNN model performed better when data from the same participant were used for training and testing. Overall, within-participant models tend to perform better than cross-participant models on decoding tasks.

**Figure 1.**
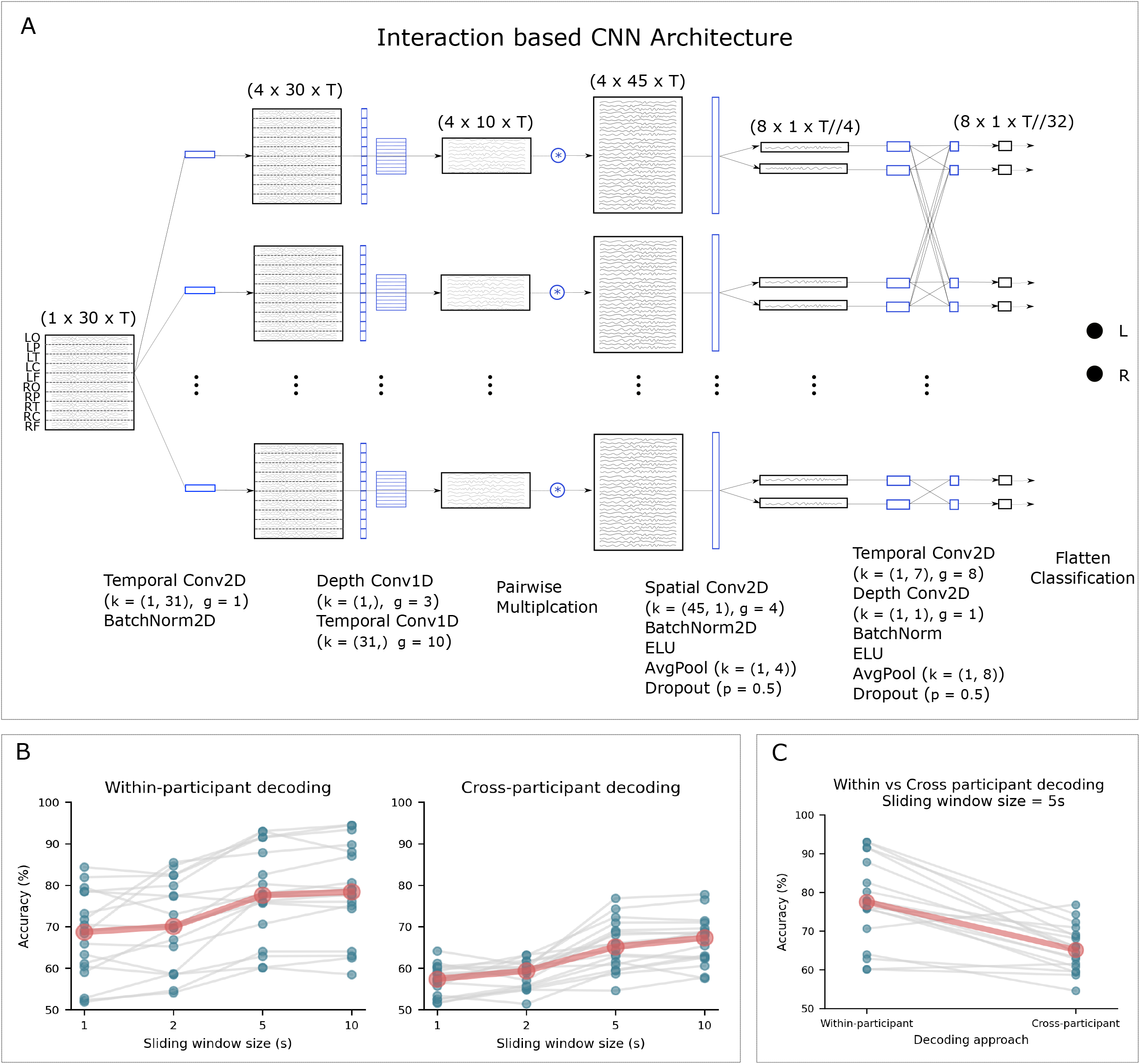
An illustration of the architecture of our CNN model and its performance on decoding spatial auditory attention within and cross participants. Panel A: The architecture of network begins with a temporal convolution, followed by a spatial convolution to reduce the dimensionality and generate a single feature time series for each cortical region. A temporal convolution is then applied, using 10 distinct kernels on each of the 10 signals. Element-wise multiplications between all pairs of the 10 time series are performed next, resulting in four matrices with a dimension of 45 x T that represent interaction time series. The subsequent convolution layer utilizes a two-dimensional spatial convolution, followed by a separable convolution comprising of a depth-wise convolution and pointwise convolutions. In the classification layer, the features are flattened and passed to a two-unit softmax classifier. The blue-colored objects in the Figure represent the CNN kernels applied at each step, while the black objects show the shape of the inputs/outputs after each step. For each convolution layer, the kernel size (denoted by k) and group parameter (denoted by g) are listed below each operator. Panel B: Within-participant (left panel) and cross participants (right panel) auditory attention decoding performance of our CNN model for four different input lengths. A sliding window of 5 s length was determined to be a fair compromise between decoding performance and data length. Blue points show per-participant results averaged over cross-validation folds. Gray lines connect the data points for single participants. Red points represent the median accuracy across participants for each input length. Panel C: Comparing within-participant and cross-participant decoding performance of our CNN model for the same participant and for the input length of 5 seconds.

### Brain interactions involved in decoding auditory attention

The architecture of our CNN model was designed to learn the interactions between ten brain regions of interest (left/right occipital, parietal, temporal, central, and frontal) and then to use only these interactions to decode the locus of auditory attention. We analyzed the spatial filters (45-dimensional vectors) obtained from the fourth convolution layer, as they give an indication of which combination of interactions between cortical regions was used by the CNN model for the decoding task. We first analyzed the CNN models obtained from the within-participant classification approach and extracted spatial filters for all cross-validations and for all participants. We next employed a two-level clustering approach where we applied k-means clustering at the individual level to reduce the data dimensionality to four spatial filters per participant. At the group level, we used hierarchical clustering to group participants with similar spatial filters into clusters. This approach yielded a total of three clusters for the spatial filters obtained from within-participant classification (see Figure 2 left panels).

The first cluster acquired from within-participant decoding comprised a spatial filter that mainly emphasizes the interactions between hemispheres in the occipital, parietal, and central regions (e.g., RO–LP, RP–LP, RP–LC, and RC–LP). These inferred interactions may be related to the lateralization of neural activity during selective auditory attention, as demonstrated by several studies [17,21,22]. In the second cluster, the spatial filter used a combination of interactions between areas located within the left hemisphere, predominantly, between the occipital, parietal, and central regions. These three cortical regions also had strong interactions with frontal areas. Indeed, this pattern of interactions between the left occipito-parietal and left frontal regions has been reported in previous studies [15,16]. In addition, the spatial filter emphasized interactions between the right parietal and left frontal areas. The spatial filter acquired from the third cluster comprised a pattern that assigned negative values for some interactions within the left hemisphere (LO–LC, LO–LP, LT–LC) and positive values for those within the right hemisphere (RO–RP, RO–RC, RP–RC, RP–RF). This spatial filter primarily uses the contrast between the interactions within the right and left hemispheres of the brain.

Analogous to the within-participants analysis, we acquired spatial filters from the cross-participant decoding approach and applied a similar clustering approach at the individual and group levels. This analysis resulted in four clusters of spatial filters (Figure 2, right panels). The first cluster obtained negative values for interactions within the left hemisphere (e.g., LO–LP, LO–LT, LP–LT, and LP–LC interactions) and positive values for those within the right hemisphere (e.g., RO–RP, RO–RT, RO–RP, and RP–RT interactions). This spatial filter utilizes the asymmetry of interactions, the difference between the interactions within the right and left hemispheres, specifically for occipital, parietal, and temporal cortical regions. In contrast, the second cluster highlights between-hemisphere interactions localized mostly to parietal and temporal regions (e.g., RT–LO, RT–LP, RT–LT, RP–LP, RP–LT). The third and fourth clusters, however, showed hemisphere-specific patterns. The spatial filter acquired from the third cluster uses mainly the interactions within the right hemisphere in the occipital, parietal, and temporal areas including e.g., RO–RC, RO–RT, RO–RP, RP–RF, RP–RC, and RP–RT. In the fourth cluster, the spatial filter uses largely the interactions within the left hemisphere between the frontal and all other regions e.g., LP–LF, LT–LF, LC–LF, LT–LC as well as some interactions within the right hemisphere including RT–RO, RT–RP, and RC-RO. These results highlight that the models trained to decode and generalize across participants relied on a combination of within-hemisphere and inter-hemisphere interactions, particularly in the occipital, parietal and temporal areas. This suggests that these cortical regions and their interactions may play a key role in facilitating selective auditory attention across individuals.

**Figure 2.**
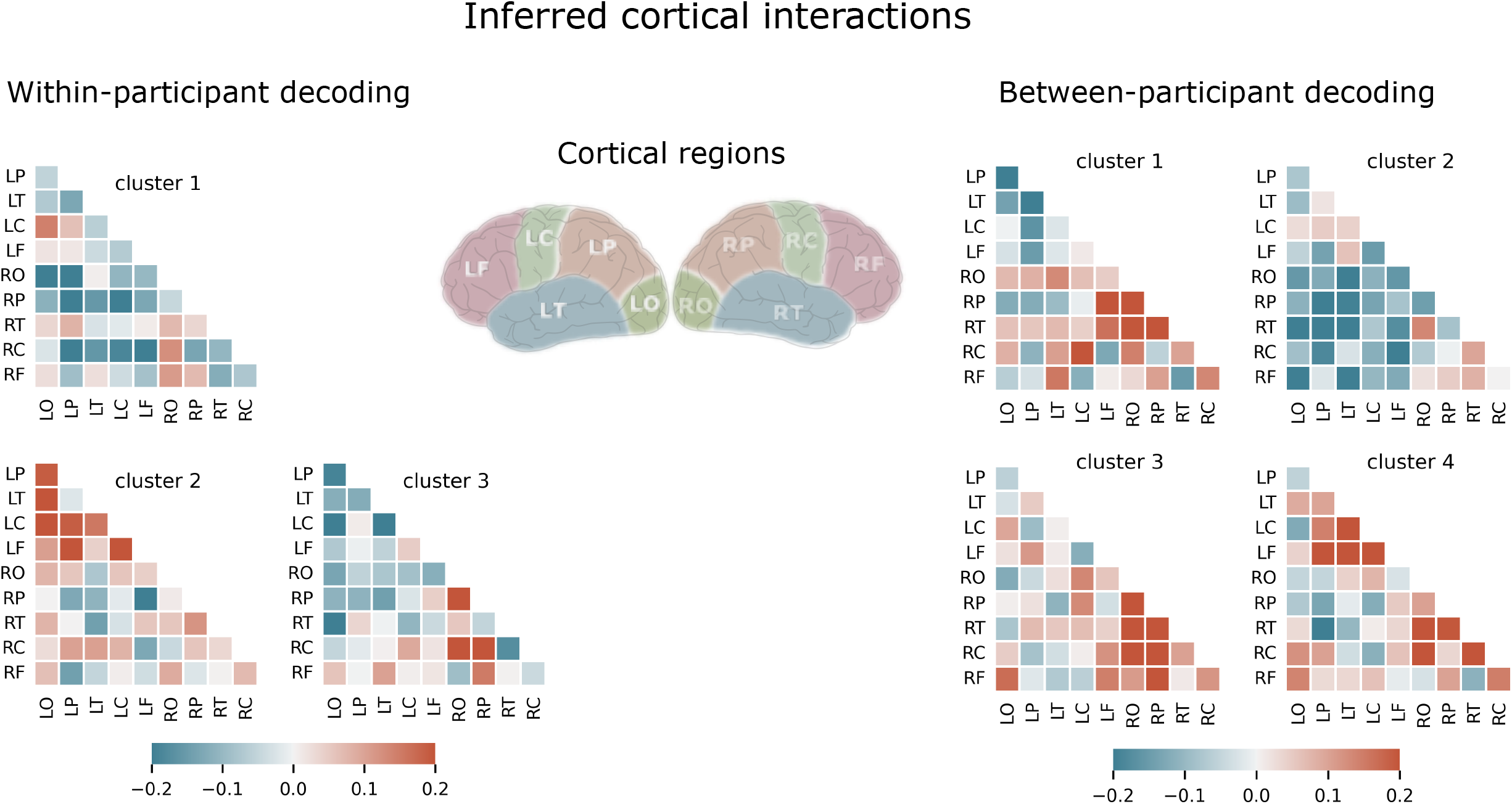
Visualization of the interactions between 10 regions of the brain derived from our CNN model trained using both within-participant (left panels) and cross-participant (right panels) decoding approaches.

### Importance of brain regions and frequency bands for spatial auditory decoding

In order to investigate the significance of the designated cortical areas for the decoding task, the architecture of our model was designed in such a way as to maintain the feature representations associated with ten cortical regions within the initial four convolution layers. This design decision allowed us to conduct a feature ablation analysis (see [28] for further details), where we eliminated the role of a specific brain region in the network by replacing the region-specific elements of the spatial filters (CNN kernels) with zeros. Specifically, we successively removed each of the brain regions (occipital, parietal, temporal, central, and frontal) from the model, and then utilized the resulting brain-region-removed model to decode the test set trials. For this analysis, we chose to remove brain regions from both hemispheres, as the removal of a single brain region from a hemisphere had a negligible impact on decoding performance. In addition to eliminating individual brain regions, we also assessed the performance of the model when brain regions within each of the left and right hemispheres were removed. The left panels of Figure 3 show the decoding performance of the model when each of the brain regions or hemispheres is removed (green boxes) as compared to the original model (orange box), for both within-participant (top panel) and cross-participant decoding (bottom panel).

During the within-participant analysis, the most significant decrease in decoding accuracy was observed upon the exclusion of the parietal (median = 72.9%, 4.66% decrease), temporal (median = 71.9%, 5.66% decrease), and central (median = 70%, 7.56% decrease) regions from both hemispheres, while the removal of the frontal (median = 78.9%, 1.34% increase) and occipital (median = 76.5%, 1.06% decrease) areas had a trivial impact on classification performance. Furthermore, omitting either full hemisphere (LH: median = 68.6%, 8.96% decrease; RH: median = 65.7%, 11.86% decrease) led to a larger reduction in decoding accuracy than omitting any single brain region alone (see Figure 3, top-left panel). We found relatively similar results for cross-participant analysis, where exclusion of parietal (median = 58%, 7.14% decrease), temporal (median = 59.1%, 6.04% decrease), and central (median = 61.5%, 3.64% decrease) regions resulted in a poor decoding performance while frontal (median = 64%, 1.14% decrease) and occipital (median = 61.9%, 3.24% decrease) regions were of lesser importance. Notably, removing the right hemisphere decreased the prediction performance, nearly to chance level (LH: median = 55.6%, 9.54% decrease; RH: median = 54.5%, 10.64% decrease).

In an attempt to discern the frequency bands employed by our CNN model in decoding auditory attention, we successively filtered out four canonical frequency bands (delta: 2-4 Hz, theta: 4-8 Hz, alpha: 8-13 Hz, and beta: 15-32) from the input data to generate band-removed inputs. Subsequently, we utilized these frequency band-removed data as inputs for the trained models obtained from both within-participant (Figure 3, top-right) and cross-participant (Figure 3, bottom-right) training approaches. The performance of our model on inputs with deleted frequency bands is depicted in green boxes, while our original findings are presented as a reference in the orange box. It is important to note that, at this stage, we only retested our previously trained models, taking care to prevent any leakage between the test and training sets. The right panels of Figure 3 demonstrate that our model’s accuracy decreased most significantly when the alpha frequencies were eliminated (within-participant: median = 70.4%; cross-participant: median = 59%), which is consistent with previous research indicating the involvement of alpha frequencies in auditory attention [17,39].

**Figure 3.**
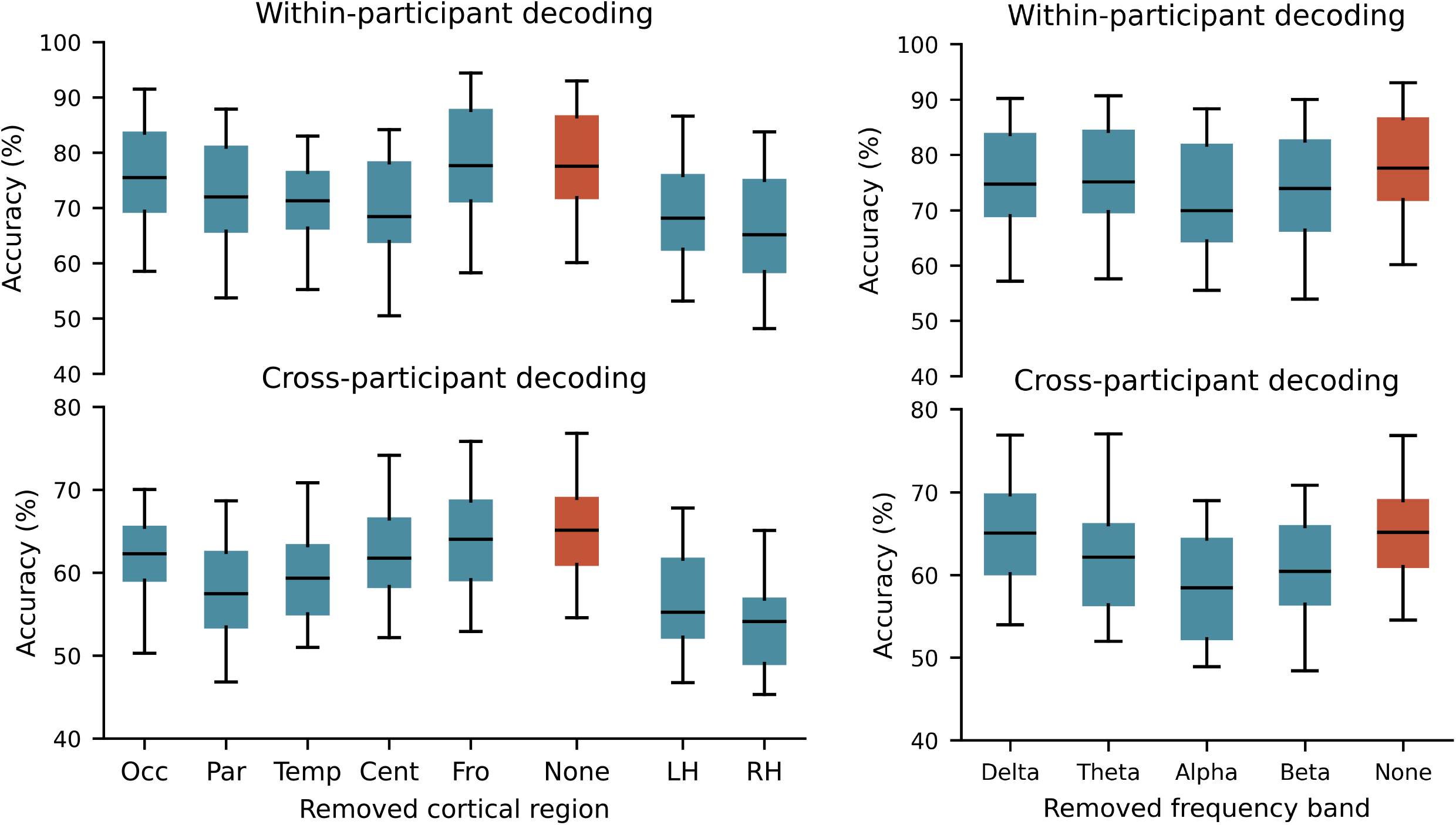
Importance of brain regions and frequency bands for decoding auditory attention. Left panels: performance of our CNN model when particular brain regions were removed from both hemispheres (first five green boxes from left), no brain region is excluded (orange box), only left hemisphere regions are removed (the second green box from right), and only right hemisphere regions are removed (the last green box). The top panel: within-participant decoding, bottom panel: cross-participant decoding. To remove brain regions from our decoding model, we set the corresponding kernels to zero (Occ: occipital, Par: parietal, Temp: temporal, Cent: central, Fro: frontal, LH: left hemisphere, RH: right hemisphere). Right panels: within-participant (top panel) and cross-participant (bottom panel) auditory attention decoding performance of the CNN model when a certain frequency band is filtered out from the input data. We retested the model (without retraining) by filtering out the delta (2-4 Hz), theta (4-8 Hz), alpha (8-13 Hz), and beta (15-32) frequency bands from test data (green boxes). The orange box shows the original results.

### Frequency specificity of identified interactions for spatial auditory decoding

Results of the analyses conducted in the previous subsections demonstrated the importance of alpha-band frequencies and the significance of interactions, primarily among the occipital, parietal, and central regions, for within-participant decoding, as well as among the parietal and temporal regions for cross-participant classification. In an effort to thoroughly explore the interrelationship between these two findings derived from independent analyses, we sought to determine whether the identified interactions are specific to a particular frequency band. To address this question, we iteratively filtered the data within the delta, theta, alpha, and beta frequency bands and constructed band-specific inputs. Additionally, we conducted a feature ablation analysis on our CNN model in which we eliminated the influence of the interactions identified through our cluster analysis. This was achieved by first Z-transforming the values within each spatial filter relative to the mean and standard deviation of that filter, then converting each Z-value to a p-value using the normal cumulative distribution function and retaining only those values with p < 0.05 (uncorrected). The ablated model was created by zeroing out the elements determined to be significant interactions in the fourth convolution layer. Finally, we tested both the original and ablated models with the band-filtered inputs without re-training. Figure 4 illustrates the decoding performance of the original model (dark-colored boxes) and ablated model (light-colored boxes) for four filtered inputs (green boxes) and for broad-band inputs (orange boxes) using within-participant and cross-participant decoding approaches. Here, the ablated models for both within-participant and cross-participant decoding approaches were created by removing the significant interactions determined from their respective first clusters.

Our analysis of the within-participant decoding approach demonstrated that the removal of significant interactions from the full model resulted in a notably decreased accuracy of decoding for alpha band data, with a median accuracy of 73.40% in the full model and 63.87% in the ablated model. This effect was less pronounced for beta band inputs, with a median accuracy of 65.25% in the full model and 61.47% in the ablated model. In contrast, the performance for delta and theta band inputs showed only minor changes, with median accuracies of 56.07% and 54.45% in the full models and 54.74% and 53.14% in the ablated models, respectively. For the cross-participant decoding approach, we observed a large decrease in performance in the ablated models for both alpha-and beta-band inputs, with median accuracies of 59.66% and 60.36% in the full models and 56.41% and 54.68% in the ablated models, respectively. In contrast, the delta-and theta-band inputs showed only trivial decreases, with median accuracies of 52.12% and 56.90% in the full models and 52.13% and 54.03% in the ablated models, respectively. Our results show that the interactions determined by the first cluster for within-participant decoding are primarily dominated by alpha band frequencies, while for cross-participant decoding, both alpha and beta frequencies contribute. Figure S2 represents the results for all clusters determined by our cluster analysis for both within-participant and cross-participant decoding methods, showing that almost for all clusters the elimination of corresponding significant interactions led to a large decrease for alpha and beta band inputs.

**Figure 4.**
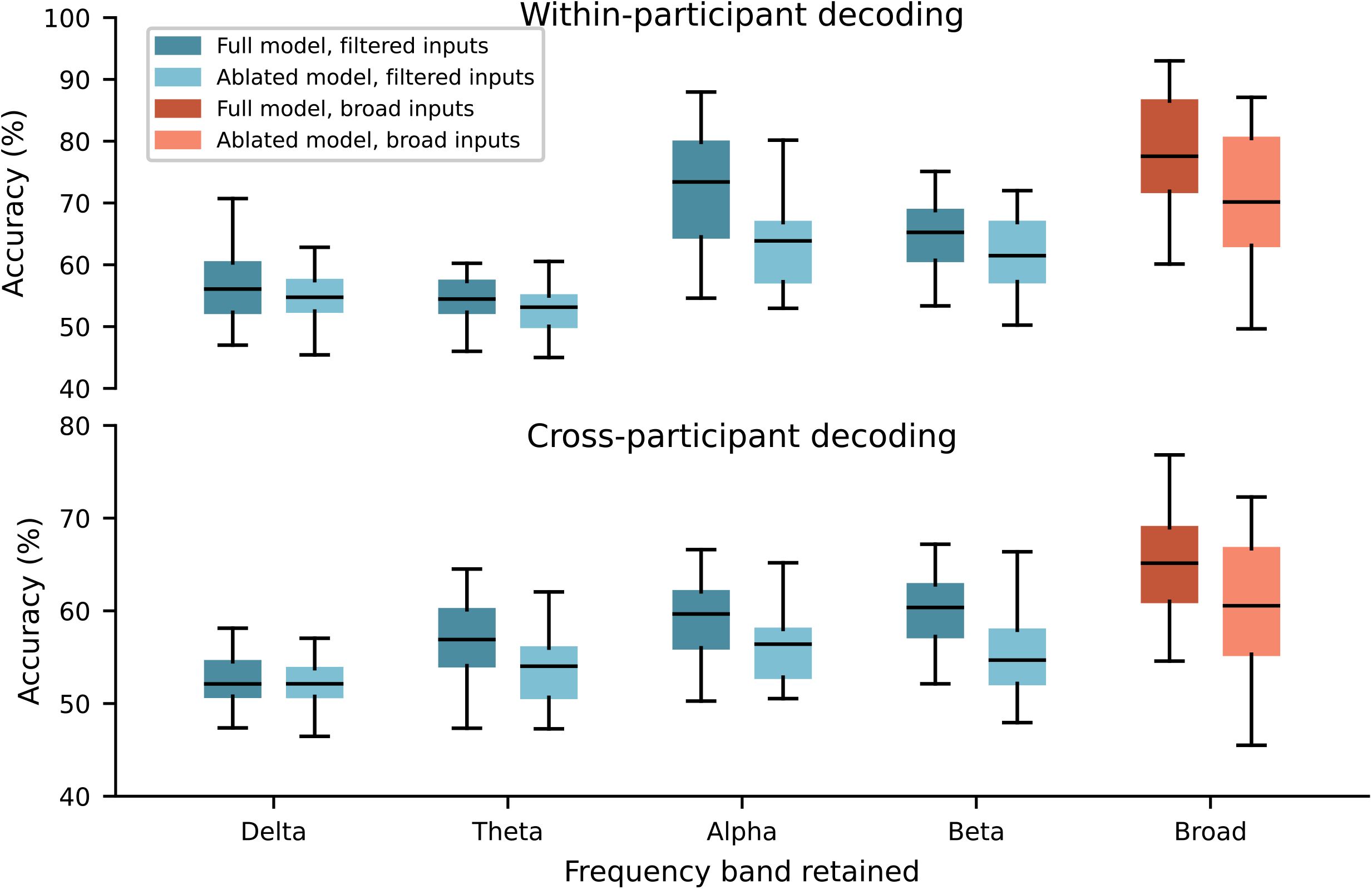
Performance of our convolutional neural network model when data filtered within four frequency bands (delta, theta, alpha, and beta) were used as inputs. Both the original CNN model (dark-colored boxes) and an ablated model (light-colored boxes) were tested using within-participant (top panel) and cross-participant (bottom panel) decoding approaches. Results for broad-band (2-32 Hz) inputs are shown in orange boxes. The ablated model was constructed by removing interactions identified as significant by the first cluster."

## Discussion

In this study, we utilized source-reconstructed and anatomically-resolved EEG data as inputs for a novel CNN topology that we propose as a new analysis tool for enhancing our understanding of the significance of cortical interactions in decoding auditory spatial attention. Our CNN model was specifically designed to learn pairwise interaction representations for 10 cortical regions including the left and the right occipital, parietal, temporal, central, and frontal areas. Using only these features for decoding, our model achieved median accuracy of 77.56% for within-participant classification and 65.14% for cross-participant.

Upon analyzing the spatial filters learned by the CNN model, we uncovered the presence of three main types of interactions utilized by the model for the decoding task: inter-hemisphere interactions, hemisphere-specific interactions, and the contrast between interactions within the right and left hemispheres. The inter-hemisphere interactions were localized in the occipital, parietal, and central regions for the within-participant decoding approach, and in the parietal, temporal, and frontal regions for the cross-participant decoding approach. Further ablation analysis revealed that these interactions were dominated by alpha frequencies in both decoding approaches. Hemisphere-specific interactions were localized only in the left hemisphere for within-participant decoding, but in the right hemisphere or a combination of both hemispheres for cross-participant decoding. Our analysis also demonstrated that these interactions were largely dominated by alpha frequencies, with some contribution from beta frequencies. Finally, the pattern of interactions encapsulating the difference between interactions within the right and left hemispheres primarily utilized interactions between parietal and central regions for within-participant decoding, and interactions between the parietal and temporal regions for cross-participant decoding. Ablation analysis showed that both alpha and beta frequencies contributed to these interactions.

These findings, obtained from our analysis approach, reflect the results of previous EEG studies demonstrating the presence of at least two distinct generators of alpha oscillations over central and parieto-occipital regions [17,21,22] and lateralization of parietal alpha activity as the main indicator of spatial attention [17–20]. In addition, our results are further supported by a recent study that applied a CNN model to sensor-space EEG data and uncovered the involvement of beta-band activations, primarily in the frontal and temporal regions, in the decoding of spatial auditory attention [14]. Our confirmatory results demonstrate the utility of our analysis approach, which employs a CNN architecture built based on the characteristics of anatomically resolved EEG data, in utilizing features known to be important in auditory attention. As a result, this strategy can be utilized as an analysis tool to deduce additional characteristics of brain functioning related to auditory attention.

Although our findings highlight the crucial role of alpha and beta frequencies in decoding spatial attention, we observed a minimal effect of lower frequencies, such as delta and theta bands. This concurs with recent research reporting the trivial impact of low frequencies on auditory attention [14]. However, it is possible that this may also be due to the architecture of the CNN model, which may not be sufficiently deep to capture longer structures in the data. In the context of deep neural networks applied to computer vision, it is well-established that initial convolution layers capture simple structures, while deeper layers build upon one another and learn to encode more abstract structures in the data [40]. However, implementing a deeper CNN would lead to an increase in the number of parameters and, in our case, overfitting due to the relatively small number of trials per participant and a small overall number of participants. Other factors that can affect the frequency of inputs include the size of convolution kernels and the use of pooling layers. In our study, in order to supply the interaction operator with a broad range of frequencies, we set the size of convolution kernels to be half of the sampling frequency, allowing the convolution to capture all frequencies above 2 Hz and included average pooling layers only after the interaction layer. Alternative network architectures that may be more effective in capturing longer temporal structures like delta and theta frequencies in the data include those designed for language modeling such as recurrent neural networks or convolution-based approaches like WaveNet [23,41].

In the present study, in line with several studies which have attempted to adapt existing CNN architectures to the properties of EEG data rather than importing them directly from the computer vision applications [27,28,35,37], we sought to utilize CNNs not merely as a high-performing "black box" decoder, but rather as an interpretable tool for deriving task-specific interactions between brain regions. To achieve this, we first transformed the input data into an interpretable structure by source projecting the EEG data and obtaining time series for individual cortical regions of interest. Additionally, we integrated the shape of the input data within the architecture of the model by maintaining the anatomical information in the initial four convolution layers. This way, the model was pushed to leverage the existing prior domain knowledge and subsequently utilize it for the decoding task. This input and architecture integrated approach allowed us extract task-specific interactions from trained models.

Although the median accuracy of our decoding pipeline was 77.56% for within-participant and 65.14% for cross-participant training approaches, it varied amongst participants for both within-participant and cross-participant training approaches. For example, as shown in Figure 1, for some participants the decoding performance was better than 90%, whereas for two participants it was around chance level. This discrepancy may be related to the relatively low number of trials per participant for the within-participant approach and the small number of participants for cross-participant generalization. However, we were unable to determine why certain individuals had low performance in our decoding process. To properly account for this inter-individual heterogeneity, particularly for real-time applications like hearing aids, future research may use more data, employ data augmentation techniques, utilize more flexible CNN architectures, and/or include behavioral data.

The methodology applied in this study to learn brain interactions from data and then infer the learned interactions from trained models can be generalized to other EEG and MEG experiments with different decoding tasks by applying minor adjustments to the architecture of our CNN model.

In conclusion, we presented a CNN that takes in time series data from 10 cortical regions and learns to extract the relevant interactions between them for decoding auditory spatial attention. Our interpretable model design is based on the properties of EEG time series data. By interpreting the trained models, we found that the network was able to identify known important interactions in spatial auditory attention and that alpha and beta frequencies played a key role in its performance. Overall, our CNN approach provides a promising approach for exploring and understanding neural dynamics and their interactions involved in decoding tasks.

## Materials and Methods

### Dataset

We used publicly available EEG data recorded from 18 participants with normal hearing. EEG was recorded from 64 channels (BioSemi ActiveTwo system) with sampling rate of 512 Hz. Two additional electrodes were also placed on the mastoids as physiological reference signals [42]. During the experiment, sixty 50-s long trials were recorded, during which participants listened to two simultaneously presented speech streams (one on the left and one on the right) and were cued to listen to one and ignore the other. Virtual auditory environments (VAEs) were simulated using the room acoustic modeling software Odeon. The binaural VAEs were reproduced in a soundproof, electrically-shielded listening booth with ER-2 insert earphones (Etymotic Research). To spatially separate speech signals presented via earphones, the speech signals were convolved with non-individualized head-related impulse responses for azimuth angles of ±60° and an elevation angle of 0°. The order of presentation of different virtual auditory environments (anechoic, mild reverberation, and high reverberation) was independently randomized across trials for each participant. Moreover, the position of the target speaker relative to that of the listener (±60°) as well as the gender of the target speaker were randomized across blocks for each participant. The presentation order of the stories was also randomized across participants. For further details about the data and experiment see [42].

### EEG preprocessing

We used the MNE-Python package [43,44] to analyze the data. EEG time series were band-pass filtered using a zero-phase forward filter with range 2–32 Hz and down-sampled to 64 Hz. EEG data were then epoched based on the provided trigger information, and we discarded trials where only a single speaker was presented. Next, the time series of each channel was visually inspected and excessively noisy channels/time segments were manually rejected. Data were re-referenced to the common average of channels. Infomax-based independent component analysis [45] was applied to identify and remove artifactual components from the EEG recordings (average: 4.7, std: 2.5). Retained components were back-projected to the sensor space.

### EEG source analysis

To reconstruct time courses of the cortical sources generating the scalp EEG activity, we carried out source reconstruction, which requires solving forward and inverse models. The forward model describes the physical process of neuronal current propagation from the dipolar sources constrained to cortical regions to the EEG channels. To compute the forward model, we used a standard template anatomy based on FSAverage included in the Freesurfer package (https://surfer.nmr.mgh.harvard.edu), and extracted the inner skull, outer skull, and outer scalp surfaces, each comprising 1280 nodes. These three surfaces were used to obtain a three-layer head model (conductivity of the layers was set to MNE-Python defaults: 0.3, 0.006, 0.3). The source space was defined as a dipole grid on the white matter surface down-sampled to 5124 source points using the topology of a recursively subdivided icosahedron (“ico-4” option). Finally, the forward model was computed, using the boundary element method (BEM) as implemented in the MNE-Python package, between sources constrained to the cortical surface and 64 EEG electrodes projected to the scalp surface.

Three-dimensional dipolar sources were reconstructed under free-orientation using the minimum norm estimate method [46] with depth weighting of 0.8 to compensate for the depth bias towards the superficial sources [47]. The cortical surface was parceled into 10 coarse regions-of-interest (Figure S1) including right and left occipital (RO, LO), right and left parietal (RP and LP), right and left central (RC, LC), right and left frontal (RF, LF), right and left temporal (RT, LT) regions according the Desikan-Killiany atlas [48] to obtain time courses for each region.

To reduce the number of time series per ROI, we performed dimensionality reduction using the Generalized Eigenvalue Decomposition broadband (GEDb) approach [49]. GEDb seeks components that enhance signal to noise ratio in a certain frequency band of interest by solving a generalized eigenvalue decomposition problem where the covariance of band-pass filtered data is specified as the signal covariance and the covariance of the broadband data as the noise covariance matrix (or reference covariance matrix) [49,50]. GEDb was carried out on each ROI across time courses of sources within the ROI, at delta-theta (2–8 Hz), alpha (8–13 Hz), and beta (15–32 Hz) frequency ranges, separately. Only the first component of GEDb was retained yielding three time series (corresponding to three frequency bands) per ROI and 30 time series per participant.

Next, by applying a sliding window of size T and with an overlap of 50% over time points, we generated the input samples of size 30 x T for the classifier. We tested four values for T (1 s, 2 s, 5 s, and 10 s).

Our goal in applying source reconstruction and GEDb analysis was to provide anatomically-resolved and SNR-enhanced data for our CNN model and ultimately be able to interpret our results in a neurophysiologically meaningful way.

### Convolutional neural networks

A convolutional neural network (CNN) is a type of artificial neural network that consists of a series of convolutional layers and nonlinear activation functions. These layers employ kernels or filters of a specified size, which slide over the data to extract local features. When applied to EEG data, these convolution kernels can extract either temporal or spatial features, depending on their shape. For instance, a row kernel can be used to learn temporal features, while a column kernel can be utilized to extract spatial information hidden in the raw data. Convolutional layers are often followed by pooling layers, which down-sample the feature space but retain important features. Activation functions introduce nonlinearity to the model. The CNN is optimized by minimizing a loss function using an optimization algorithm, which estimates the optimal parameters (kernels) for the model.

Our proposed CNN for decoding auditory attention is shown in Panel A of Figure 1. The architecture of this model is partly inspired by an existing CNN called EEGNet (Lawhern et al., 2018); however, it has been modified to learn interactions between 10 cortical regions and utilize only those features for the decoding task. The inputs to the model are matrices of 30 x T, where 30 represents the number of signals acquired from 10 brain regions and T denotes the number of time points within the sliding window. These signals have undergone GEDb dimension reduction to enhance activity within the delta-theta, alpha, and beta frequency bands. In the initial layer of our model, we employed a two-dimensional convolution with four row kernels of size (1, 31), yielding four feature maps of dimensions 30 x T. The length of the temporal kernel was set to 31, corresponding to half the sampling rate, thereby enabling the model to capture frequencies within the range of 2-32 Hz. This convolution layer was then succeeded by batch normalization along the time points [51].

In the subsequent layer, we utilized a one-dimensional convolution with a column kernel of size (30,) and a group parameter equal to the number of brain regions. This operation is equivalent to applying 10 distinct convolution layers, each processing signals from three different cortical regions and producing a single output signal for each region. These output signals are then concatenated to generate a 10 x T feature map. Through this operation, the model is able to learn a linear combination of the three signals for each cortical region.

We then implemented a one-dimensional temporal convolution with 10 groups of convolution kernels, each of size (31,). This operation serves to further refine the information extracted by the previous layer by enabling the model to learn more complex patterns within each of the signals, and is especially useful when the signals contain a mixture of frequencies or when the patterns of interest are not temporally synched across signals.

Next, we performed element-wise multiplication between all pairs of the 10 time series, resulting in four matrices with dimensions 45 x T that represent the interaction time series. This operation forces the model to learn the interactions between the 10 time series and use these features for the decoding task.

In the following convolution layer, we used a two-dimensional spatial convolution of size (45, 1) to learn spatial filters from the interaction time series that determine which linear combination of interaction time series is most important for the decoding task. We then applied batch normalization and the exponential linear unit (ELU) activation function [52]. To reduce the number of time points, we employed an average pooling layer of size (1, 4). The average pooling operation reduces the sampling frequency of the data to 16 Hz. The kernels utilized as spatial filters here were also constrained to maintain a maximum norm of one. After that, a dropout with a probability value of 0.5 was applied [53]. The dropout operation helps to prevent overfitting by randomly dropping out a portion of the activations in the layer during training, effectively forcing the model to learn more robust features.

The subsequent layers of our CNN model were designed similar to the architecture of EEGNet [28]. Specifically, we employed a separable convolution, which consists of a depth-wise convolution followed by pointwise convolutions [54]. Unlike standard convolution, which performs both spatial and temporal computation in a single step, separable convolution divides the computation into two distinct stages: the depth-wise convolution applies a single convolutional filter to each input channel, while the pointwise convolution combines the outputs of the depth-wise convolution via a linear operation. When applied to EEG data, this technique allows the model to learn feature maps in the time domain and subsequently combine these maps in an effective manner. In addition, separable convolution also has the advantage of reducing the number of parameters to be learned.

In the final layer, the classification layer, the inclusion of a fully connected layer was omitted in order to decrease the number of parameters and incorporate features indicating interactions across brain regions directly into the classifier. To do that, the features were flattened and passed to a two-unit softmax classifier [28,55]. The model was implemented using Pytorch (https://pytorch.org).

### Model training and evaluation

To train, evaluate, and test our model, we used an approach similar to the procedure described by Lawhern and colleagues [28], and performed within-participant and cross-participant analyses to assess the decoding of auditory attention.

For the within-participant decoder, we employed a block-wise four-fold cross validation approach, where two blocks out of a participant’s four total blocks were set as the training set, one block as the validation set, and the last block as the test set.

For the cross-participant decoder, each individual participant’s full dataset was used as the test set five different times. On each of those five folds, four participants’ data were randomly selected as the validation set, and the remaining 13 participants’ data were used as the training set. Since we repeated this five times for each of 18 participants, this resulted in 90 different folds for cross-validation.

The model was trained by minimizing the cross-entropy loss between the model outputs and the corresponding labels using Adam optimizer [56]. We set the batch size to 32 and omitted the bias term from all convolutional layers. We ran 200 training epochs with validation stopping, saving the model parameters that produced the lowest validation set loss.

### Clustering

After training the CNNs for all cross validations and individuals, we extracted the spatial filters learned in the third convolution layer for all models. We then employed k-means clustering at the individual level followed by a hierarchical clustering at the group level to partition the spatial filters into distinct clusters.

At the individual level, to reduce the data dimensionality, the spatial filters obtained from cross-validations per participant were grouped into four clusters using the k-means algorithm. The k-means algorithm treats each spatial filter as a point in 45-dimensional space and groups data into k mutually exclusive clusters by minimizing the centroid distance of observations within clusters and maximizing the distance between clusters. We used the cosine distance metric. The number of clusters was set to four, consistent with the number of spatial filters in our CNN model. The choice of four clusters for k-means was also matched with the optimal number of k-mean clusters identified by the Elbow curve and the Silhouette methods. The k-means algorithm was repeated 10 times with different (randomly determined) centroids and maximally 100 iterations. The solution with the smallest sum of distance values within the clusters was accepted.

At the group level, we employed a hierarchical clustering approach, to group participants with similar spatial patterns into subgroups. In a bottom-up manner, this algorithm initially treats each data point as a cluster of its own and then further pairs of clusters are successively merged as one moves up the hierarchy. We used cosine angle as the metric for the linkage computation. Finally, based on our visual inspection of the resulting dendrograms and local maxima of the resulting silhouette values, we chose three clusters for within-participant analysis and four clusters for the cross-participant training approach. For our clustering analyses we used the Scikit learn package (<u>scikit-learn.org</u>).

