## Supplementary information for "Convolutional neural networks can identify brain interactions involved in decoding spatial auditory attention"

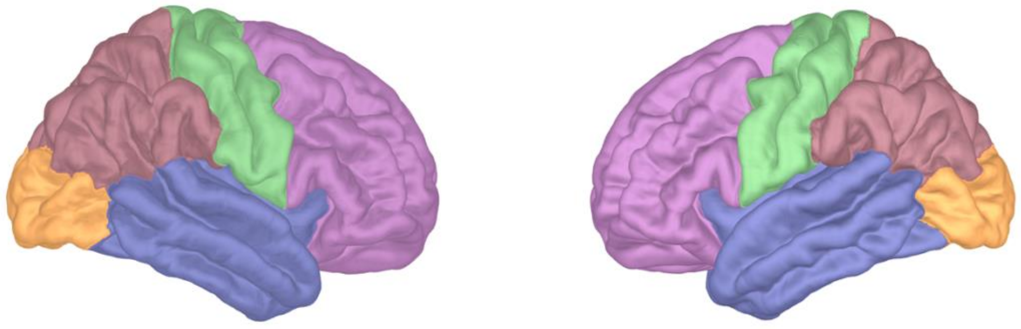

**Figure S1.** Parcellation of the cortex surface used for EEG source reconstruction including left and right frontal, central, temporal, parietal, and parietal regions.

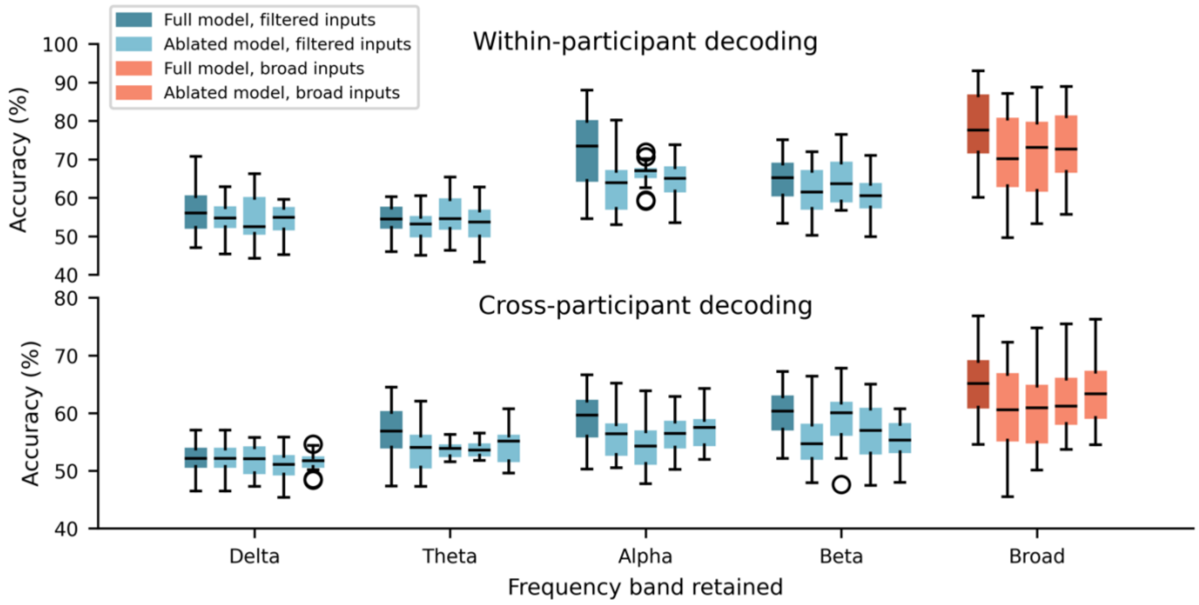


**Figure S2.** Performance of our CNN model when the data were filtered within the four frequency bands (delta, theta, alpha, and beta) and used as inputs for both original (full) CNN model (dark-colored boxes) and the ablated model (light-colored boxes) for within-participant (top panel) and cross-participant (bottom panel) decoding approaches.

Orange boxes show the results for broad-band (2-32 Hz) inputs. The ablated model was constructed by eliminating important interactions detrimined by the first (first light blue box from the left), second (second light blue box from the left), third (third light blue box from the left), and fourth (only for cross-participant decoding, fourth light blue box from the left) clusters. Outlier points were determined based on the first and third quartiles (Q1, Q3), and inter-quartile range (IQR) as values outside the range [Q1 – 1.5*IQR, Q3 + 1.5*IQR], according to default settings of the boxplot function implemented in Matplotlib package.
